# Hypoxia-inducible factor 1α (HIF1α) Suppresses Virus Replication in Human Cytomegalovirus Infection by Limiting Kynurenine Synthesis

**DOI:** 10.1101/2021.01.12.426401

**Authors:** Lisa M. Wise, Yuecheng Xi, John G. Purdy

## Abstract

Human cytomegalovirus (HCMV) replication depends on the activities of several host regulators of metabolism. Hypoxia-inducible factor 1α (HIF1α) was previously proposed to support virus replication through its metabolic regulatory function. HIF1α protein levels rise in response to HCMV infection in non-hypoxic conditions, but its effect on HCMV replication was not investigated. We addressed the role of HIF1α in HCMV replication by generating primary human cells with HIF1α knocked out using CRISPR/Cas9. When HIF1α was absent, we found that HCMV replication was enhanced, showing that HIF1α suppresses viral replication. We used untargeted metabolomics to determine if HIF1α regulates metabolite concentrations in HCMV infected cells. We discovered that in HCMV-infected cells, HIF1α suppresses intracellular and extracellular concentrations of kynurenine. HIF1α also suppressed the expression of the indoleamine 2,3-dioxygenase 1 (IDO1) rate-limiting enzyme in kynurenine synthesis. In addition to its role in tryptophan metabolism, kynurenine acts as a signaling messenger by activating aryl hydrocarbon receptor (AhR). Inhibiting AhR reduces HCMV replication while activating AhR with an exogenous ligand increases HCMV replication. Moreover, we found that feeding kynurenine to cells promotes HCMV replication. Overall, our findings indicate that HIF1α reduces HCMV replication by regulating metabolism and metabolite signaling.

**Importance:** Viruses, like human cytomegalovirus (HCMV), reprogram cellular metabolism using host metabolic regulators to support virus replication. Alternatively, in response to infection, the host can use metabolism to limit virus replication. Here, our findings show that the host uses hypoxia-inducible factor 1α (HIF1α) as a metabolic regulator to reduce HCMV replication. Further, we found that HIF1α suppresses kynurenine synthesis, a metabolite that can promote HCMV replication by signaling through the aryl hydrocarbon receptor (AhR). In infected cells, the rate-limiting enzyme in kynurenine synthesis, indoleamine 2,3-dioxygenase 1 (IDO1), is suppressed by a HIF1α-dependent mechanism. Our findings describe a functional connection between HIF1α, IDO1, and AhR that allows HIF1α to limit HCMV replication through metabolic regulation, advancing our understanding of virus-host interactions.

## Introduction

Human cytomegalovirus (HCMV) is a herpesvirus that establishes lifelong asymptomatic infection in most people. HCMV infection in people with a compromised immune system causes disease that can lead to death, and congenital infection is a leading cause of birth defects (1). Replication of HCMV depends on evading cellular innate antiviral responses and hijacking host processes to support virus replication. Infection alters activity in many pathways in the host metabolic network, such as increasing glycolysis and the flow of carbons into lipid synthesis to support virus replication (2–8).

HCMV metabolic reprogramming involves hijacking the activity of host metabolic regulators (2). Hypoxia-inducible factor 1α (HIF1α) is a metabolic regulator that is altered by HCMV infection. Cell sensing of HCMV infection increases HIF1α protein levels under conditions with normal oxygen levels (i.e., normoxia) (9). Infected cells sustain HIF1α activity upon expression of HCMV early genes (10). These previous works proposed that HIF1α was induced by HCMV to support virus replication and pathogenesis (9, 10). However, the biological significance of HIF1α in HCMV infection remains unaddressed.

Here, we examined the role of HIF1α in HCMV replication by infecting CRISPR/Cas9 engineered HIF1α knockout (KO) primary human cells in normoxia. In contrast to the proposal of the previous work, we found that HCMV replication is enhanced in HIF1α KO cells. This observation suggests that HIF1α activity in infected cells alters metabolism as a protective strategy to limit viral infection. Some metabolites or metabolic activities support antiviral response (11). We determined if HIF1α regulation of metabolism is necessary for it to reduce HCMV replication using untargeted metabolomics. Among the metabolites regulated by HIF1α in HCMV-infected cells was kynurenine (KYN). KYN intracellular and extracellular levels were markedly increased in HCMV-infected cells lacking HIF1α. KYN is a metabolite in tryptophan degradation that is made by the indoleamine 2,3-dioxygenase 1 (IDO1) pathway. We show that blocking IDO1 reduced HCMV replication, suggesting that KYN synthesis supports HCMV replication. In addition to its metabolic function, KYN also acts as a signaling messenger by activating the aryl hydrocarbon receptor (AhR). We found that blocking AhR signaling reduced HCMV replication, suggesting that HIF1α imparts an antiviral effect by decreasing KYN synthesis and AhR activation in HCMV infection. Overall, our findings demonstrate that HIF1α in HCMV infected cells under normoxic conditions regulates metabolism in a way that reduces virus replication through a functional connection between HIF1α, IDO1, and AhR.

## Results

### HIF1α reduces HCMV replication

HCMV infection using the AD169 or Towne lab-adapted strains increases HIF1α protein in the presence of oxygen (9). We expanded on this observation by determining if the HIF1α protein levels are affected by infection with the TB40/E strain that has not been extensively passaged in fibroblasts. In TB40/E-infected cells at a multiplicity of infection (MOI) of 3 infectious units per cell, HIF1α protein levels are increased by >3-fold in the early stage of HCMV infection (Figure 1A-B). Like the lab-adapted strains, HIF1α protein levels were highest at 1 day post-infection (dpi) in cells infected with TB40/E (9, 10). In our experiments, the cells were mock-infected or infected when they were subconfluent and maintained in normoxic conditions throughout the timecourse. In these conditions, the uninfected cells continued to grow, and HIF1α levels rose throughout the 4 d timecourse (Figure 1A). This observation was expected since hypoxic microenvironments created in cells after reaching full confluence increase HIF1α protein levels (12, 13). In contrast, HIF1α levels rose and then decreased in the HCMV-infected cells (Figure 1A). Next, we examined if HCMV infection alters HIF1α activity by measuring the expression of VEGF, a gene that is transcriptionally regulated by HIF1α. We found that VEGF transcripts were 20-fold higher in HCMV-infected cells than mock-infected cells, indicating that HIF1α is active in HCMV-infected cells maintained in normal oxygen conditions (Figure S1A).

**Figure 1.**
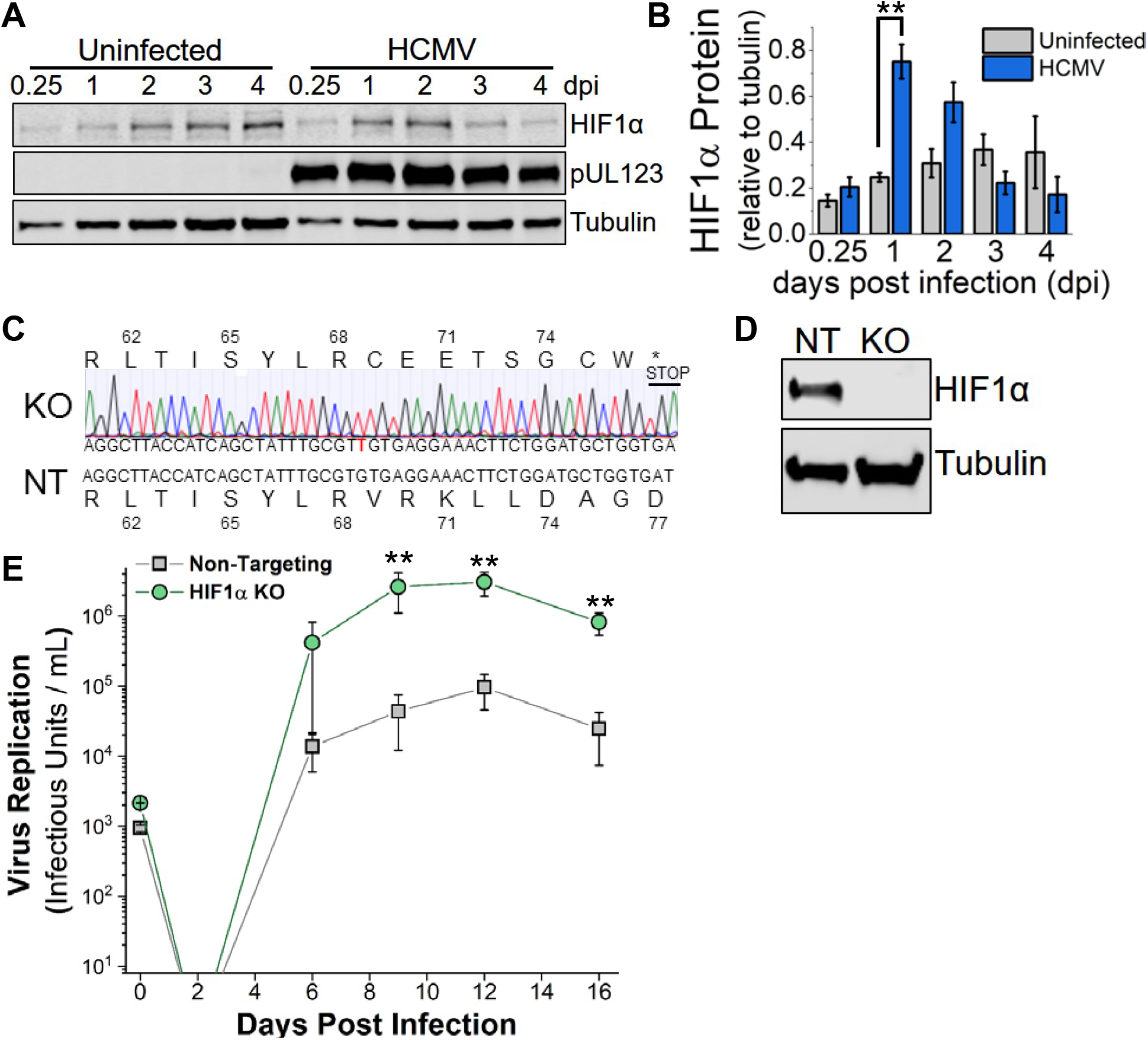
HIF1α is induced by HCMV infection and limits virus replication in normoxia. **(A)** HIF1α protein levels in uninfected and HCMV-infected fibroblasts cells were determined by Western blot. Cells were infected at MOI=3 infectious units per cell with TB40/E strain when cells were 80% confluent. Uninfected cells continued to grow, reaching full confluency by day 1-2. **(B)** Quantification of HIF1α protein relative to tubulin in cells described in A. Data are represented as mean ± standard deviation (S.D.), and significance was determined by unpaired t test, N=3 (**p<0.01). **(C)** HIF1α KO cells were generated using CRISPR/Cas9 in HFF-hTERT cells. Shown is the Indel sequencing for HIF1α KO and the sequence from cells treated with a non-targeting (NT) gRNA. **(D)** HIF1a protein level in NT and HIF1α KO cells following 24 h treatment with 100 μM CoCl2 to mimic hypoxia. **(E)** HIF1α KO and NT cells were infected at a MOI=0.05, and virus growth was measured by tissue culture infectious dose 50 (TCID50) at the indicated days post infection (dpi). Data are represented as mean ± S.D., and significance was determined by ANOVA, Tukey Test, N=3 (**p<0.01) (day 6 excludes an outlier, which is shown in Figure S1).

We generated HIF1α knockouts (KO) in primary human fibroblasts using CRISPR/Cas9 to determine the effect of HIF1α on HCMV replication. We identified HIF1α KO by sequencing (Figure 1C). Our HIF1α KO clone contained a single nucleotide insertion that caused a frameshift that resulted in a premature stop codon. We confirmed the loss of HIF1α protein by Western blot. We treated cells with cobalt chloride (CoCl2) to chemically mimic hypoxia and increase HIF1α protein levels. Consistent with our sequencing results, we observed no HIF1α protein in the HIF1α KO cells (Figure 1D). As a control, we generated CRISPR/Cas9 cells expressing a non-targeting (NT) gRNA that does not recognize any human or HCMV gene. Next, we examined the ability of HCMV to replicate in HIF1α KO and NT cells. We infected HIF1α KO and NT control cells at a low MOI of 0.05 infectious unit per cell and quantified the production of new virus progeny over 16 days. Starting at 9 dpi, HIF1α KO cells produced ~60-fold more infectious HCMV progeny than NT cells (Figure 1E and S1B). At 12 and 16 dpi, ~30-fold more infectious virus was produced by HIF1α KO cells than NT cells, suggesting that HIF1α suppresses HCMV replication and may contribute to an innate cellular antiviral response.

### Metabolic regulation by HIF1α in HCMV-infected cells

Since HIF1α regulates metabolism (14–16), we investigated if HIF1α controls a metabolic barrier to HCMV replication. We used an untargeted liquid-chromatography high-resolution tandem mass spectrometry (LC-MS/MS) metabolomic approach to determine if HIF1α regulates metabolism in HCMV-infected cells. In these experiments, we measured intracellular and extracellular metabolites extracted from infected HIF1α KO and NT cells in normoxic conditions at 2 dpi. We selected 2 dpi since HIF1α protein levels are greatest at early time points post-infection (Figure 1A), HIF1α activity was high at 2 dpi as measured by VEGF transcript levels (Figure S1A), and significant metabolic reprogramming by the virus occurs at 2 dpi (3, 4, 17). First, we analyzed the untargeted data set against a library of metabolites from several metabolic pathways, including glycolysis, TCA cycle, amino acids, and nucleotide metabolism. The library was built using commercially purchased metabolite standards that we used to define LC retention times and MS/MS spectral features using our LC-MS/MS methods. These defined retention times and MS/MS features were searched our untargeted data to find peaks that matched these defined (i.e., ‘known’) metabolites. Of the approximately 50 metabolites examined using this method, the abundance of most were unaltered in the HCMV-infected HIF1α KO cells compared to infected NT control cells (Figure S2A-B). Of the eight metabolites altered by HIF1α KO, four are involved in nucleotide metabolism, and three are intermediates in glycolysis or TCA cycle.

Next, we used the metabolomic analysis software, MAVEN (24), to select mass spectral peaks unbiasedly (Table S1). MS1 peaks were selected using a high-resolution setting of ≤5 parts per million (ppm). We filtered the data to determine peaks of significant interest for identification (Figure 2A). In these filtering steps, we removed peaks caused by contaminants (i.e., background noise). We determined contaminants using two methods. First, potential contaminants from the buffers used in LC, the HPLC columns, and mass spectrometer were defined. Second, we identified potential contaminants from our extraction method. In each metabolomic experiment, cells were seeded in a six-well plate where at least one well contained no cells. The ‘no cell’ control was handled in parallel with the cells throughout the entire extraction and LC-MS/MS processes. Using these methods to identify contaminants, we retained MS peaks for further analysis only if they had a ≥10:1 signal-to-background ratio. Since MS/MS information is needed to confirm metabolite identification, the data was further filtered to retain MS1 peaks that had associated MS/MS fragments. Next, peaks were retained if observed in both the HIF1α KO and NT samples for ≥3 independent experiments. The peaks that passed our filtering process were used to determine statistical significance between HIF1α KO and NT samples and organized by fold change in HIF1α KO relative to NT (Figure 2B).

**Figure 2.**
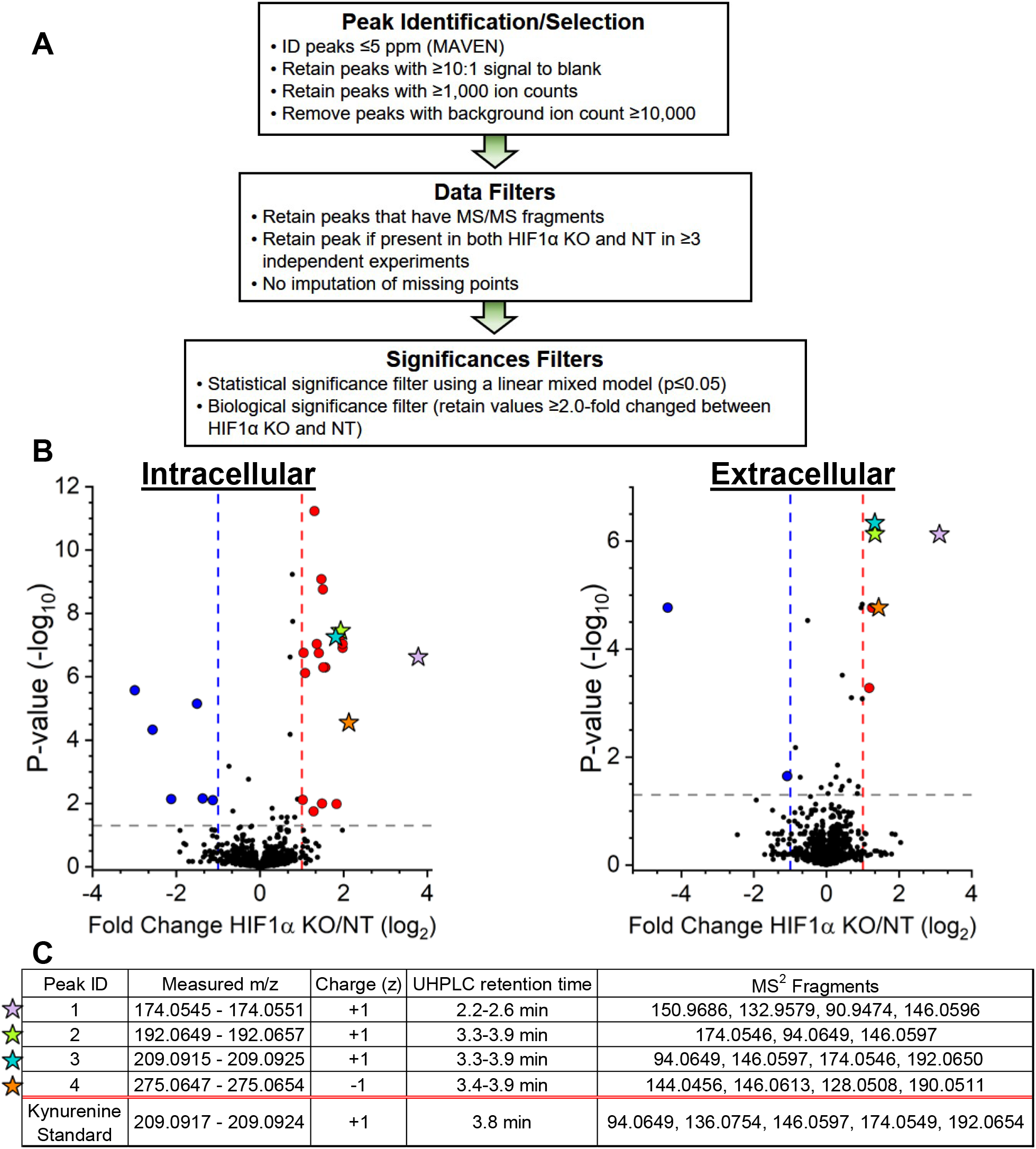
Untargeted metabolomics of HCMV-infected HIF1α-KO and non-targeting (NT) control cells. **(A)** Schematic of the workflow for analyzing untargeted metabolomics LC-MS/MS data. Mass spectral peaks were selected and quantified using (MAVEN) software. Peaks were defined using a high mass accuracy setting of less than 5 parts per million (ppm). Peaks were then selected if they had ≥1,000 ions to ensure quality of quantitation above the limit of detection and the presence of MS/MS fragments that can be used to confirm metabolite identification. Finally, statistical significance was determined using linear mixed-effects models as described in the SI Materials and Methods Details. A ≥2-fold change between HIF1α KO and NT was used to highlight potential biological significance. **(B)** At 2 dpi, intracellular and extracellular metabolites were extracted from HIF1α KO and NT cells infected with TB40/E at a MOI = 3. Metabolite peaks that passed the selection and data filters described in part A are presented. Metabolites that decrease (blue) or increased (red) by ≥2-fold in HIF1α KO cells relative to NT cells are shown as colored circles or stars. Stars represent four metabolites that were increased in both the intracellular and extracellular environments of HIF1α KO relative to NT cells. The data is from 3-6 independent experiments. **(C)** Mass spectral and HPLC information of the four metabolites starred in part B. Also shown is information obtained using a commercially produced kynurenine standard that was analyzed using LC-MS/MS conditions used for untargeted metabolomic.

The abundance of most intracellular metabolites was similar in HIF1α KO and NT cells (Figure 2B). We observed a ≥2-fold increase in 20 peaks and ≥2-fold decrease in 6 peaks in HIF1α KO cells relative to NT cells. Fewer extracellular metabolites—including fewer metabolites that were HIF1α-dependent—were observed than in the intracellular fraction (Figure 2B). We found two peaks that were ≥2-fold decrease in HIF1α KO cells relative to NT cells. None of these peaks matched the peaks in the intracellular fraction that were decreased by HIF1α KO.

In the extracellular fraction, we observed six peaks increased by HIF1α KO (Figure 2B). Notably, we four of these peaks were increased in both the intracellular and extracellular fractions by ≥2.5-fold in HIF1α KO cells relative to NT control cells (Figure 2B peaks represented by stars). We prioritized the identification of these four peaks. Peaks 2 and 3 had matching retention times of 3.3-3.9 min in reverse-phase LC conditions and shared several MS/MS fragments (i.e., 94.06, 146.05, and 174.05) (Figure 2C). The data for these peaks were collected over a nine-month period that involved several batches of LC buffers and columns that resulted in 0.6 min shift in retention time. The MS1 m/z of peak 3—209.09—matches the calculated m/z of kynurenine (KYN) within our defined 5 ppm range. We used a commercially obtained KYN standard to confirm its MS1 m/z and determine its retention time using our reverse-phase LC method and its MS/MS fragments using our metabolomic methods. The retention time and MS1 m/z for the KYN standard matched those we observed for peak 3 (Figure 2C). Next, we examined the MS/MS fragments. The four MS/MS fragments of peak 3 matched four of the top five fragments we observed using the KYN standard (Figure 2C), confirming the identification of peak 3 as KYN.

When defining the MS1 m/z of KYN in positive mode using the commercial KYN standard, we observed that electrospray ionization was fragmenting KYN to form an ion at 192.06 m/z. This ionization fragment matched the measured MS1 m/z for peak 2 within 5 ppm. The three MS/MS fragments observed for peak 2 matched three fragments observed using the KYN standard and fragments observed in peak 3 (Figure 2C), providing evidence that both peaks represent KYN. We conclude that peak 2 and 3 are KYN. Peaks 1 and 4 remain unidentified. Since the levels of KYN in human serum were previously found to be associated with HCMV infection (18, 19), we decided to focus on the role of KYN in virus replication.

### HCMV-infection raises KYN levels, which are suppressed by HIF1α

We further investigated the relationship between HIF1α and KYN levels. In uninfected cells growing in normoxia, the loss of HIF1α has little to no effect on intracellular or extracellular KYN levels (Figure 3A-B). In cells, the NT cells that express HIF1α, HCMV-infected cells have a 3.5-fold increase in intracellular KYN than uninfected cells at 2 dpi (Figure 3A). Similarly, KYN levels were greater in the extracellular fraction of HCMV-infected NT cells compared to uninfected NT cells (Figure 3B). This data demonstrates that HCMV infection, in cells with HIF1α, enhances intracellular and extracellular KYN levels.

**Figure 3.**
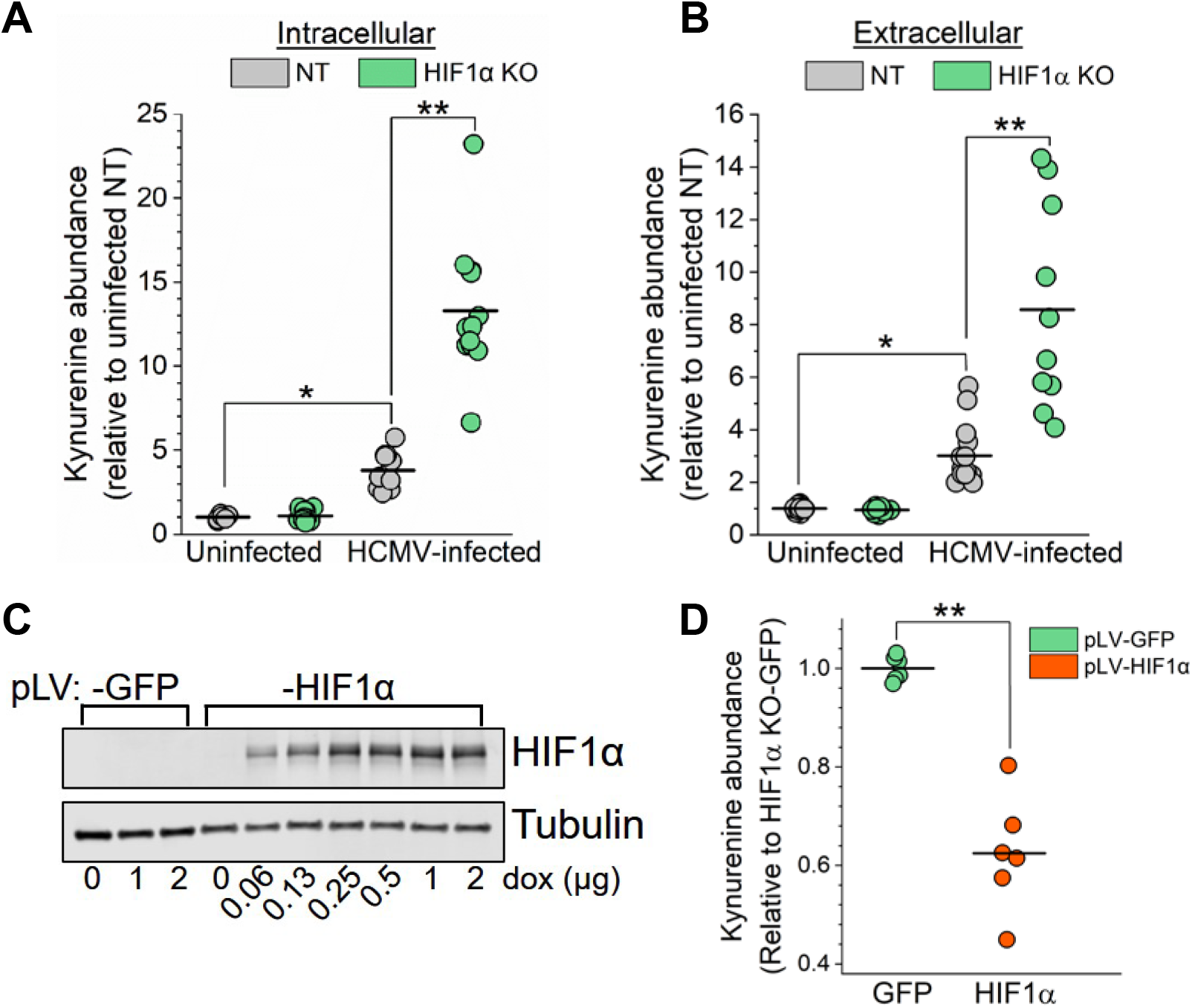
HIF1α suppresses kynurenine (KYN) levels in HCMV-infected cells. **(A)** Intracellular KYN levels in uninfected and HCMV-infected NT and HIF1α KO cells were measured by LC-MS/MS. Cells were infected at MOI=3, and metabolites were extracted at 2 dpi. The mean is represented by a black bar, and significance was determined by 2-way ANOVA, Tukey Test, N≥6. (*p<0.05; **p<0.01). **(B)** KYN levels in the extracellular environment in conditions described in part A. The mean is represented by a black bar, and significance was determined by 2-way ANOVA, Tukey Test, N≥6. (*p<0.05; **p<0.01). **(C)** Western blot analysis of HIF1α KO cells expressing GFP or HIF1α containing a silent mutation to mutate the Cas9 PAM recognition site. GFP and HIF1α expression was induced using a doxycycline-inducible system. Cells were treated with increasing concentrations of doxycycline for 2 d. **(D)** Intracellular KYN levels in HCMV-infected HIF1α KO cells expressing GFP or HIF1α-Cas9 PAM mutant. Cells were infected for 1 h at MOI=2 and then treated with 1 μg/mL doxycycline prior to extraction of metabolites at 2 dpi. The mean is represented by a black bar, and significance was determined using an unpaired t test, N=3 (*p<0.05; **p<0.01).

Relative to HCMV-infected NT cells, infected HIF1α KO cells had 3.5-fold and 2.5-fold higher intracellular and extracellular KYN levels (Figure 3A-B). Next, we sought to confirm that KYN is regulated by a HIF1α-dependent mechanism using a second independently-engineered CRISPR/Cas9 HIF1α KO clone that contained a different gRNA. We generated a second HIF1α KO clone with two deletions that introduced frameshift mutations and failed to express HIF1α protein (Figure S3A-B). At 2 dpi, KYN levels in HIF1α KO clone 1 and clone 2 were ≥2-fold higher than in NT cells, further supporting the conclusion that KYN levels are regulated by HIF1α (Figure S3C).

For further confirmation that HIF1α suppresses KYN levels in HCMV-infected cells, we re-expressed HIF1α in our KO cells using a doxycycline-inducible system (Figure 3C). In these cells, we engineered the re-expressed HIF1α to contain a silent mutation that removes the Cas9 PAM recognition site while leaving the amino acid sequence of the protein unaffected. As a control, we expressed green fluorescent protein (GFP) in HIF1α KO cells. We infected these cells for 1 hpi, then washed and treated the cells with doxycycline to induce HIF1α or GFP expression. KO cells re-expressing HIF1α had almost 2-fold lower levels of KYN than HIF1α KO cells expressing GFP (Figure 3D). These observations provide further evidence that HIF1α suppresses KYN levels in HCMV-infected cells.

### IDO1 and aryl hydrocarbon receptor (AhR) activities promote HCMV replication

Since KYN is elevated in HCMV-infected HIF1α KO cells, we determined if the expression of the rate-limiting enzyme in KYN synthesis, indoleamine 2,3-dioxygenase 1 (IDO1), is regulated by a HIF1α-dependent mechanism. At 2 dpi, IDO1 transcripts were increased by >2-fold in HCMV-infected HIF1α KO cells relative to infected NT cells (Figure 4A). Our observations that IDO1 transcripts, KYN levels, and HCMV replication are enhanced in HIF1α KO cells lead us to hypothesize that IDO1 activity promotes HCMV replication. We tested HCMV replication is dependent on IDO1 activity in HIF1α KO cells using the IDO1 inhibitor NLG919. First, we determined if NLG919 treatment at 100 nM was cytotoxic. At 9 days post-treatment, cell survival in DMSO and NLG919 treated cells were equivalent (Figure S4). We infected cells using the same conditions for the virus growth assay comparing HIF1α KO and NT cells (Figure 1E). At 9 dpi, infectious HCMV progeny production was >2-fold reduced in cells treated with 100 nM NLG919 than DMSO-treated cells (Figure 4B). We conclude that IDO1 activity promotes HCMV replication in HIF1α KO cells.

**Figure 4.**
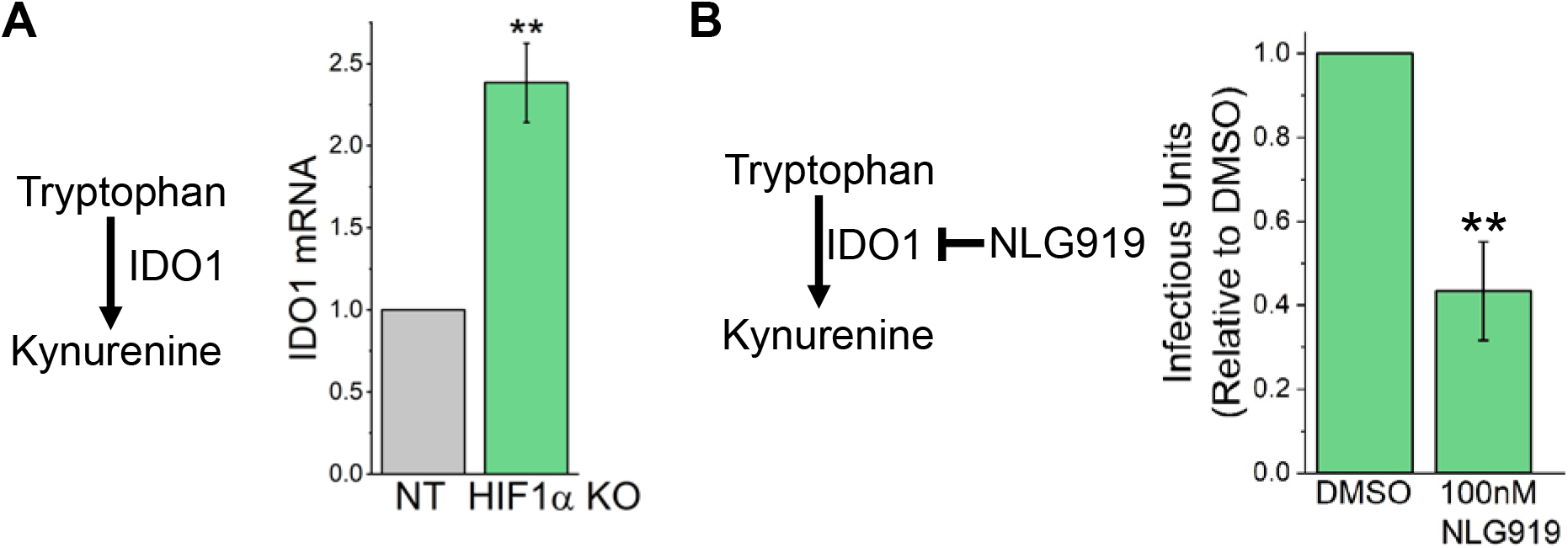
HIF1α suppresses IDO1 expression, which supports HCMV replication in HIF1α KO cells. **(A)** IDO1 is the rate-limiting enzyme in KYN synthesis. IDO1 transcripts in HCMV-infected HIF1α KO and NT cells were measured at 2 dpi using RT-qPCR. Cells were infected at MOI=3 with TB40/E. The data are represented as mean ± S.D. relative to IDO1 levels in HCMV-infected NT cells. Statistical significance was determined using an unpaired t test, N=3 (**p<0.01). **(B)** HCMV replication in HIF1α KO cells treated with an IDO1 inhibitor, NLG919, was determined. Cells were infected at MOI=0.05, and infectious progeny virus was determined at 9 dpi. At 3 and 6 dpi, the growth medium was replaced to replenish NLG919. The data are represented as mean ± S.D. relative to DMSO, and significance was determined by unpaired t test, N=3 (**p<0.01).

KYN is a ligand that activates the transcriptional activity of aryl hydrocarbon receptor (AhR). We tested if AhR is required for HCMV replication using CH223191, an AhR inhibitor that blocks KYN binding (20, 21). At 9 d post-treatment, DMSO-treated cells and 3 μM CH223191-treated cells had the same level of cell survival (Figure S4). At 9 dpi, HIF1α KO and NT cells treated with CH223191 produced fewer infectious progeny than those treated with DMSO (Figure 5A). Conversely, activation of AhR with an exogenous dioxin ligand enhances HCMV replication (22). We tested if AhR activation by an exogenous ligand would enhance HCMV replication by treating cells with 2,3,7,8-tetrachlorodibenzo-p-dioxin (TCDD) AhR activator. Since TCDD is a liquid at room temperature, we diluted TCDD directly in the cell growth medium. As a control, we add an equal volume of water to growth medium. At 9 d post-treatment, cells survived equally in water and 3.1 nM TCDD treatment (Figure S4). In NT cells, HCMV infectious progeny production was 2.5-fold higher in TCDD-treated cells than water-treated cells (Figure 5B). In HIF1α KO cells, virus progeny production was 1.5-fold higher in TCDD-treated cells than water-treated cells, demonstrating that TCDD treatment had a large effect on HCMV replication in NT cells than HIF1α KO cells (Figure 5B). Overall, these findings show that AhR activity supports HCMV replication.

**Figure 5.**
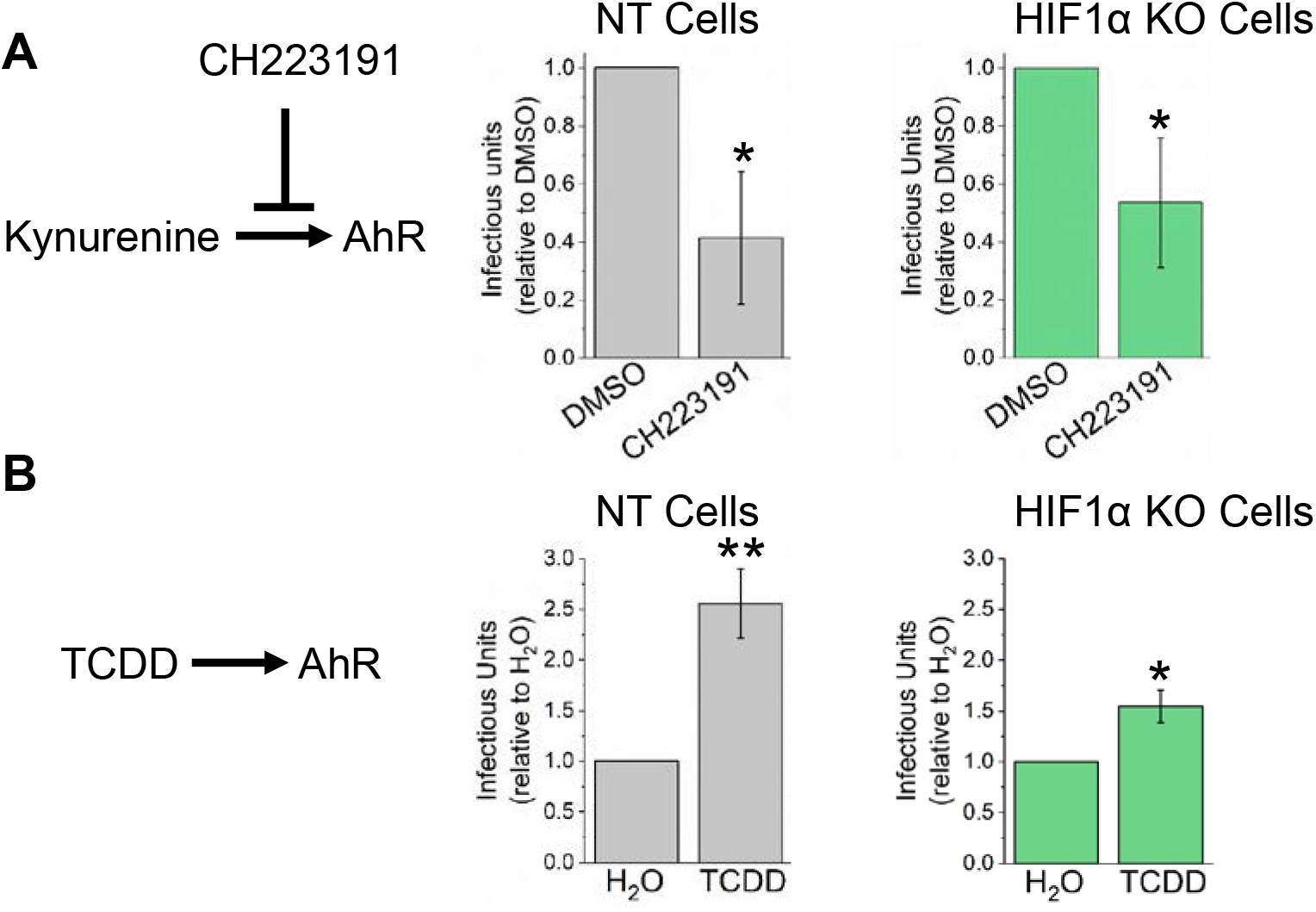
KYN receptor aryl hydrocarbon receptor (AhR) promotes HCMV replication. **(A)** HCMV replication was measured in cells treated with 3 μM CH223191, an AhR inhibitor. Cells were infected at MOI=0.05, and infectious HCMV progeny was measured at 9 dpi. At 3 and 6 dpi, the growth medium was replaced to replenish CH223191. The data are represented as mean ± S.D. relative to DMSO-treated. Significance was determined using an unpaired t test, N=3 (*p<0.05). **(B)** HCMV replication in cells treated with 3.1 nM 2,3,7,8-tetrachlorodibenzo-p-dioxin (TCDD), an exogenous AhR activator. Cells were infected at MOI=0.05, and infectious HCMV progeny was measured at 9 dpi. At 3 and 6 dpi, the growth medium was replaced to replenish TCDD. Since TCDD is liquid at room temperature and was directly diluted into cell growth medium, tissue culture grade water was used as a control. The data are represented as mean ± S.D. relative to water-treated. Significance was determined using an unpaired t test, N=3 (*p<0.05; **p<0.01).

### KYN enhances HCMV Replication

Based on our observations, we hypothesize that KYN enhances HCMV replication. We tested this hypothesis by feeding KYN to cells following a low MOI infection and monitoring HCMV spread in cell culture. In this experiment, we used normal primary human fibroblast cells that have not been genetically altered by CRISPR/Cas9 or any other means. At 1 hpi, we washed the cells and then fed the cells growth medium supplemented with 0, 0.25, 0.5, or 1.0 mM KYN. At 6 dpi, we fixed and visualized infected cells using an antibody against HCMV IE1 protein (Figure 6A). Since IE1 is localized in the nucleus, we determined the number of IE1-positive nuclei per plaque. We counted the number of infected cells per plaque in 6 independent experiments for a total of ≥80 plaques for each KYN concentration tested. In untreated cells, the average plaque size was 10 cells per plaque. In KYN treated cells, the average plaque size was 15-23 cells (Figure 6B). Since KYN treatment increased HCMV plaque size, we conclude that KYN promotes HCMV infection.

**Figure 6.**
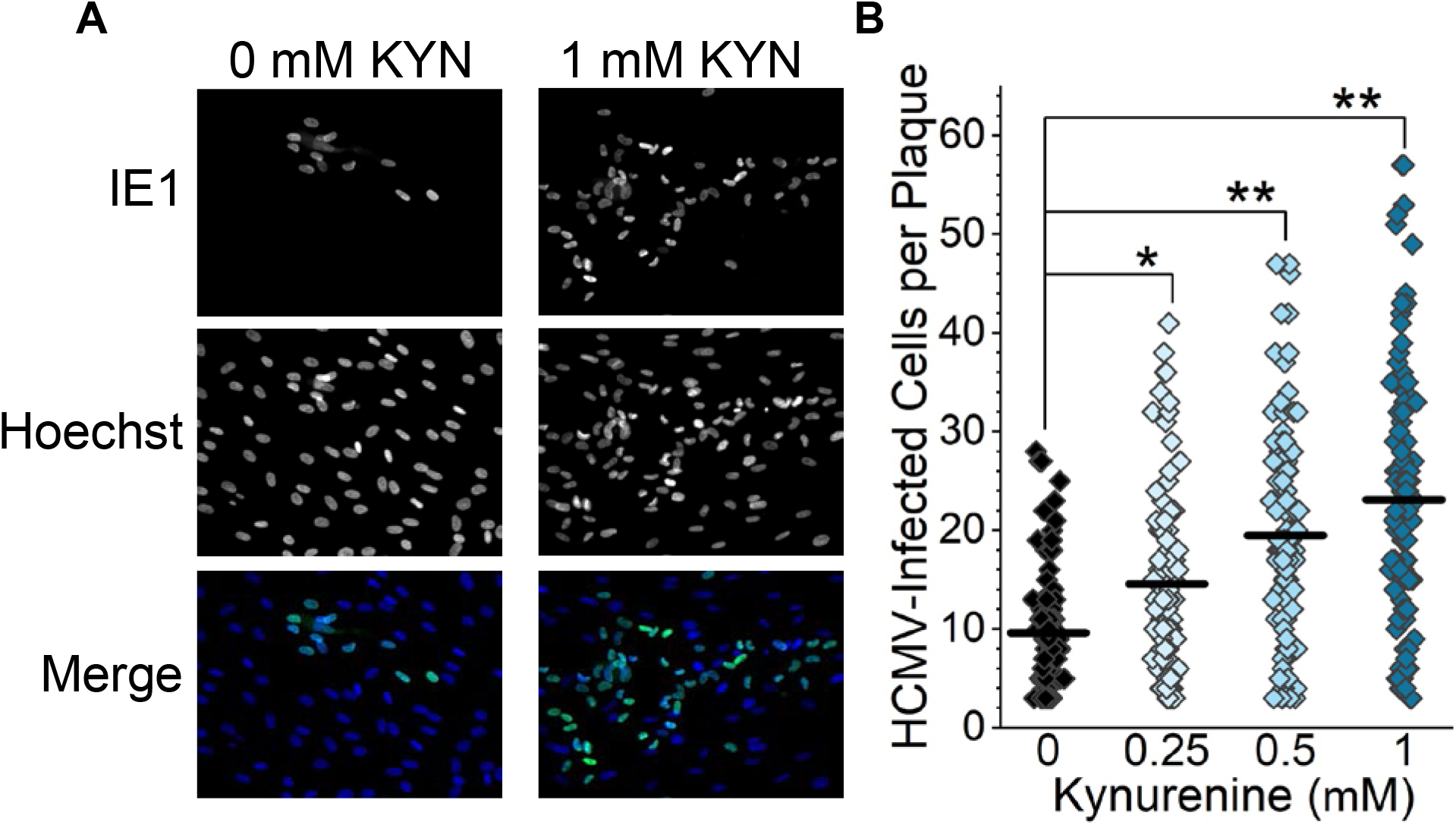
Feeding KYN to cells promotes HCMV infection. **(A)** HCMV replication and spread in fibroblast cells treated with KYN was measured. Cells were infected with 100 infectious units for 1 h and then washed with PBS. The infected cells were then overlaid with growth medium containing 0, 0.25, 0.5, and 1 mM KYN and 0.6% methylcellulose. At 3 dpi, the growth medium was replaced to replenish KYN. At 6 dpi, the cells were washed and fixed with methanol. The nuclei of infected cells were visualized using immunofluorescence and an antibody against HCMV IE1 protein. All nuclei were visualized by staining with Hoechst 33342. Representative images from cells treated with 0 and 1mM KYN are shown. (B) The number of IE1 positive cells per plaque were counted in cells treated with KYN as described in part A. At least 10 plaques were counted per experiment. In order to be counted, a plaque must have contained at least three IE1 positive cells. The mean is represented as a black line, and significance was determined by Kruskal-Wallis ANOVA and Mann-Whitney test, N=6 (*p<0.05; **p<0.01).

## Discussion

Viral reprogramming of host metabolism is essential for HCMV replication (2, 23–25). Accordingly, HCMV infection reprograms the flow of nutrients through metabolic pathways by altering the activity of host metabolic regulators (5, 7, 8, 26). However, most studies have focused on how host metabolic regulation supports virus replication. Here, our findings show that the host uses HIF1α as a metabolic regulator to reduce virus replication.

McFarlane and colleagues demonstrated that UV-inactivated HCMV virus particles triggered HIF1α expression in cells in normoxia (9), consistent with the idea that the host activates HIF1α to contribute to an antiviral response. Our work extends this concept by demonstrating that IDO1 expression, KYN levels, and virus replication is suppressed via a HIF1α-dependent mechanism in HCMV-infected cells in the presence of oxygen. IDO1 is an interferon-γ inducible gene. Interferon-γ induced IDO1 gene expression and protein activity are reduced by HCMV infection (27–31). Our findings suggest that this observed reduction of IDO1 activity by HCMV infection could be due to a HIF1α-dependent host response to suppress HCMV replication. Moreover, hypoxia represses the induction of IDO1 in uninfected cells (32, 33), providing further evidence supporting the role of HIF1α in regulating IDO1 activity.

Our findings suggest that HIF1α suppressed IDO1 expression and KYN synthesis to reduce HCMV replication. Based on our observations, we propose that the host cell uses a HIF1α-dependent response to reduce IDO1 activity and limit KYN levels to dampen AhR signaling and reduce HCMV replication. In the absence of HIF1α, HCMV infection enhances IDO1 synthesis of KYN. Subsequently, KYN metabolite-mediated activation of AhR signaling would promote HCMV replication. Since we observed an increase in both intracellular and extracellular KYN levels in HIF1α KO cells, KYN activation of AhR may occur within infected cells and neighboring uninfected cells. We observed that TCDD activation of AhR promoted HCMV replication (Figure 5B). According to our proposed model, TCDD activation of AhR would be lower in HIF1α KO since those cells would have a higher level of AhR activate by KYN. Indeed, we observed that TCDD treatment promoted HCMV replication to a lesser extent in HIF1α KO cells than NT cells (Figure 5B).

In this study, we found that HCMV infection raises the intracellular and extracellular concentration of KYN (Figure 3A-B). Immunosuppressed HCMV-infected transplant recipients have elevated KYN serum levels (18). Additionally, people with HCMV and human immunodeficiency virus type 1 (HIV-1) co-infection have enhanced KYN serum levels relative to those with HIV and herpes simplex virus type 1 (HSV-1) co-infection (19). While our observations demonstrate that HCMV-infection increases KYN levels in cell culture, it remains unknown if elevated KYN serum levels *in vivo* are altered due to the metabolic activity of HCMV-infected cells or uninfected cells responding to infection. KYN functions as a metabolite in tryptophan catabolism and as a mediator in AhR signaling. Blocking AhR activity limits HCMV replication in HIF1α KO and control cells (Figure 5A). Conversely, activation of AhR enhances HCMV replication (Figure 5B) (22). While additional studies are needed, these observations suggest that elevated KYN levels in human serum may promote infection.

We found that the levels of several metabolites are HIF1α-dependent (Figure 2B). The identification of KYN allowed us to define a mechanistic connection between HIF1α, IDO1, and AhR during HCMV infection. Several additional metabolites regulated by HIF1α remain unknown in our untargeted metabolomics data (Figure 2). These include two peaks—peak 1 and 4—that are higher in both the intracellular and extracellular fraction of HIF1α KO cells relative to NT Cells (Figure 3C). These still unidentified metabolites may also regulate HCMV replication, and their future identification may result in new understandings of the interaction of HCMV and host metabolism. Some of them, like KYN, may be reduced by HIF1α to limit HCMV replication.

Overall, we discovered a metabolic pathway regulated by HIF1α in HCMV infection in a way that suppressed virus replication. These findings suggest that HIF1α acts as an antiviral factor through its activity as a metabolic regulator. HIF1α or hypoxia has been linked to innate defenses against other viruses (34, 35), parasites (32, 36), and bacteria (37), suggesting that HIF1α regulation of metabolism may have broad impacts on microbial infections. Our work described here provides a foundation for future work to understand how HIF1α may modulate metabolism to support innate cellular antimicrobial defenses.

## Materials and Methods

### Cells and Virus Infection and Virus Replication

All experiments were performed in normoxic conditions using HCMV TB40/E strain or TB40/E containing GFP (TB40/E-GFP) was used in this study (38, 39). Virus stocks were grown from fibroblast cells electroporated with a bacterial artificial chromosome containing the TB40/E-GFP genome (BAC4-TB40/E-GFP), which was kindly provided by Dr. Thomas Shenk (Princeton University). Infectious HCMV was purified and concentrated from the supernatant of infected cells by ultracentrifugation through 20% sorbitol. Viruses were tittered using tissue-culture infectious dose 50 (TCID50). Human foreskin fibroblast (HFF) cells were grown and maintained in DMEM containing 4.5g/L glucose (Gibco 11965-092), supplemented with 10% Fetal Bovine Serum (FBS; Sigma-Aldrich #12303C), penicillin and streptomycin, and 5 mM HEPES pH 7.4 (Gibco 15630-080).

HCMV TB40/E-GFP was used for all infections. One day prior to infection or mock-infection (i.e., uninfected samples), 1.6×10^5^ cells per well were seeded on 6-well. Cells were either mock-infected or HCMV-infected for 1-2 h at the indicated multiplicity of infection (MOI). After mock or HCMV infection, cells were washed with PBS and provided DMEM growth medium containing 10% FBS.

Virus replication assays were performed in RPMI 1640 (Gibco 11875-093) supplemented with 10% FBS, penicillin, streptomycin, and 5 mM HEPES. Following infection, cells were washed with PBS, and 1.5 mL of RPMI 1640 growth medium was added to each well. Samples were collected at 0, 2, 5, 9, 12, or 16 days post infection (dpi). Viruses were harvested by scraping the well and collecting the entire volume of cells and media. Samples were stored at −80°C. After thawing, each sample was briefly sonicated to release cell-associated viruses. Finally, the volume of each sample was measured, and the amount of infectious virus per mL was determined using TCID_50_.

### Protein Analysis by Western Blot

SDS-PAGE was performed using Bio-Rad anyKD and 4-20% gradient gels. Proteins were transferred to nitrocellulose membrane (Li-Cor 926-31092), and blots were blocked using 3% milk in a Tris-buffered saline containing 0.05-0.1% tween-20. The following antibodies were used to detect proteins: anti-tubulin (Sigma-Aldrich T6199), anti-IE1 (1B12, (40)), anti-HIF1α (BD Biosciences 610959), anti-mouse dylight 800 (ThermoFisher SA5-35521), and anti-rabbit dylight 680 (ThermoFisher 35568). Anti-HIF1α antibody was incubated with blocked membranes at a 1:1000 dilution for 14-16 h at 4º C. All other antibodies were incubated for 1 h at room temperature. Proteins were visualized and quantified using a Li-Cor Odyssey CLx imager and Image Studio Software. For experiments using cobalt chloride (CoCl2) to enhance HIF1α levels, cells were treated with a final concentration of 200 μM CoCl2 for 24 h prior to lysing cells for Western blot.

### Engineering of HIF1α-KO cells using CRISPR/Cas9

sgRNA sequences specific for the human gene HIF1α were designed using crispr.mit.edu and further selected for low off-target potential for both the human and HCMV genome using CasOFFinder (41). sgRNA sequences used in this study were: CRISPR gRNA HIF1α c. 1 (5’-CCATCAGCTATTTGCGTGTG-3’) and CRISPR gRNA HIF1α c. 2 (5’-TGTGAGTTCGCATCTTGATA-3’). sgRNA were cloned into LentiCRISPR-v2, a gift from Feng Zhang (Addgene 52961) (42). For the engineering of non-targeting (NT) control cells, the control gRNA (5’-CGCTTCCGCGGCCCGTTCAA-3’) sequence from the human GeCKO v2 CRISPR Knockout Library was cloned into the LentiCRISPR-v2 plasmid (42, 43).

HFF cells were used for generating HIF1α KO clones. First, human telomerase (hTERT) was exogenously expressed using pBABE-neo-hTERT (Addgene 1774, a gift from Bob Weinberg ((44)). HFF-hTERT cells were transduced with LentiCRISPR-v2 lentiviruses that carried the gRNA and Cas9. Transduced HFF-hTERT cells were selected using puromycin (Corning 61-385-RA). Next, selected cells were diluted to one cell per well on a 96-well plate and supplemented with 150 non-transduced HFF cells. Once cells reached 90% confluence in most wells, puromycin was added to selected against the non-transduced HFF cells. Surviving cells were grown up, and potential HIF1α KO clones were initially identified by Western blotting for HIF1α expression following a 24h treatment with CoCl2. The loss of HIF1α was further confirmed by sequencing using the Guide-it Indel kit (Takara #631444) and primers targeting the corresponding genomic regions. The primers used were Indel Guid-IT sequencing primer 1 (5’-CACCTGCTTCCGACAGGTTT-3’) and primer 2 (5’-GGAAACACCTGCTTCCGACA-3’). Each clone was sequenced at least 20 times to ensure biallelic mutations in HIF1α gene.

### Engineering of HIF1α KO cells to re-express HIF1α protein

HIF1α containing silent mutations in the Cas9 PAM recognition site was used to re-express HIF1α protein in HIF1α KO cells. The wildtype human HIF1α gene was obtained from pcDNA3-HIF1α plasmid (Addgene 18949, a gift from William Kaelin (45)). A “G” to “A” mutation at position 210 using site-directed mutagenesis removed the PAM recognition site for HIF1α KO clone 1 while maintaining the wildtype protein sequence of Arg at amino acid position 70. Next, the HIF1α PAM-mutant gene was inserted into pLVX-EF1a using a Takara In-Fusion HD Cloning Plus kit. Finally, HIF1α PAM-mutant gene was transferred into a pLV-TRE-blasticidin from VectorBuilder using BamHI and XbaI to generate a plasmid termed pLV-TRE-HIF1α_G210A-blast. pLV-TRE is a lentivirus system for doxycycline-inducible expression of proteins in mammalian cells. pLV-TRE-HIF1α_G210A-blast was sequenced to confirm the PAM-site mutation and that the gene would encode for wildtype HIF1α protein. For the engineering of HIF1α KO cells that expressed either HIF1α PAM mutant or GFP as a control, HIF1α KO clone 1 cells were first treated with lentivirus particles from 293 T transfected with pLV-rtTA-hygro. Next, HIF1α KO-rtTA cells were treated with lentivirus particles generated in 293T cells transfected with pLV-TRE-HIF1α_G210A-blast or pLV-TRE-GFP_blast. All pseudo-lentiviruses were produced using pMD.2G and psPAX2, as previously described (5). Doxycycline was used to induce the expression of HIF1α or GFP in HIF1α KO clone 1 cells. For KYN measurements in HIF1α KO cells expressing HIF1α or GFP, the cells were HCMV infected for 1 h, washed, and then provided fresh growth medium containing 1 μg doxycycline. GFP expression was confirmed using a fluorescent microscope, and HIF1α expression was measured by Western blot.

### Metabolite Extraction

For all metabolomic experiments, duplicate samples were extracted in parallel for each condition. For extraction of extracellular metabolites, 150 μL of growth media was harvested from plates containing either mock-infected or HCMV-infected. The extracellular growth media was centrifuged 21,000x g for 5 min to pellet cell debris. Next, 100 μL of the resulting supernatant was used for metabolite extraction. Metabolites were extracted by adding cold methanol for a final concentration of 80% methanol. Subsequently, the samples were incubated on dry ice or at −80° C for 10-15 minutes and centrifuged at 4,000x g at 4° C to remove cell debris and proteins. Finally, the extracted metabolites were dried under nitrogen gas.

Intracellular metabolites were methanol extracted as previously described (3, 26). First, the growth media was removed, and cells were quickly washed with PBS. Next, cold 80% methanol was added to quench all metabolic reactions. Metabolites were extracted following incubation on dry ice or at −80° C for 10-15 minutes and centrifuged at 4,000x g at 4° C. Extracted metabolites were dried under nitrogen.

We identified possible contaminants using controls that lacked cells. In this case, growth medium was placed in a 6-well plate that lacked cells and analyzed in parallel to samples from 6-well plates that contained cells. The LC-MS/MS signal from the “no cell” samples was used to define the background to remove contaminates from our datasets.

### Metabolomics

All metabolites were identified and quantified using liquid chromatography tandem mass spectrometry (LC-MS/MS). Extracellular metabolites were normalized by extraction volume. Intracellular metabolites were normalized by cell volume (17, 26, 46). Metabolites were analyzed by reverse-phase or HILIC chromatography. For reverse-phase analysis, metabolites were resuspended in MS-grade water. For HILIC analysis, metabolites were resuspended in 1:1 MS-grade water and MS-grade methanol. All samples were analyzed within 24 h of extraction to limit metabolite degradation. Reverse phase analysis was performed using a Kinetex 1.7 μm F5 UPLC column (Phenomenex 00F-4722-AN), while HILIC analysis used an Acquity BEH HILILC 1.7 μm column (Waters 186003462). All LC was performed using a Vanquish LC system (Thermo Fisher Scientific) using an autosampler that stored samples at 4° C and a temperature-controlled column holder that kept columns at 25° C. Two buffers were used for LC, 97:3 water:acetonitrile plus 0.1% formic acid (buffer A) and 100% acetonitrile (buffer B). Each LC run was 30 min using the following conditions: 0% B for 2 min, 10% for 3 min, hold at 10% B for 1 min, 20% B for 4 min, hold at 20% B for 1 min, 55% B for 7 min, hold at 55% B for 1 min, 96% B for 4 min, and hold at 96% B for 1.5 min (reverse phase), and 95% B for 2.5 min, 40% B for 8 min, hold at 40% B for 7.5 min, and 95% B for 2.5 min (HILIC). All LC was performed at 0.25 mL/min, and the column was equilibrated between each sample. Untargeted metabolomics was performed using a Q-Exactive Plus orbitrap mass spectrometer (Thermo Fisher Scientific). MS1 data was collected by full scans from 60 to 900 m/z using a mass resolution setting of 140,000. Data-dependent MS/MS was performed using a TopN setting of 5. Additional settings used were: 1e^6^ AGC, 250 ms maximum injection time. Normalized collision energy (nce) was set at 40 or 45. MS2 data was collected using a resolution setting of 35,000 with 1e^5^ AGC, 120 ms maximum injection time, and 10s dynamic exclusion. Metabolites were ionized using a heated electrospray ionization probe with the following settings: sheath gas flow 20, auxiliary gas flow 10, sweep gas flow 1, spray voltage 2.5, capillary temperature 243, S-lens RF 65, and auxiliary gas temperature 205 for negative mode, and: sheath gas flow 32, auxiliary gas flow 10, sweep gas flow 1, spray voltage 3.0, capillary temperature 253, S-lens RF 58, and auxiliary gas temperature 180 for positive mode. Metabolites were analyzed using MAVEN (47, 48). UPLC- or MS-grade solvents were used for LC-MS/MS analysis, including Optima water (Fisher W7-4), Optima LC/MS methanol (Fisher A456), Optima LC/MS acetonitrile (Fisher A955), and Optima LC/MS-grade formic acid (Fisher A117).

For untargeted analysis, peaks were selected using the following settings in MAVEN: 5 ppm mass resolution, 10 scans time resolution, 5 minimum signal-to-baseline ratio, 5 minimum signal-to-blank ratio, 3 scan minimum peak width, and 2,000 ions minimum peak intensity. Background signal from samples that contained no cells was used to remove peaks from contaminates. Peaks from multiple replicates were combined using hierarchical clustering (Ward’s method) of m/z and retention time. Peaks must have been present in three of six independent experiments to be included in further analysis. Peaks were further analyzed, as outlined in Figure 2A. Briefly, peaks were retained for analysis if the total ion count was ≥1,000 ions, and if they had a ≥10:1 signal-to-blank ratio. Additionally, peaks with a background signal ≥10,000 ions were removed. Next, peaks with collected MS2 fragments were retained, while those without MS2 fragments were removed. MS/MS identification involved integration of MS2 data using a Python script (https://github.com/lisawise/maven_explorer). Finally, statistical analysis was performed on the remaining peaks, as described below.

Metabolite identification was based on three parameters: the retention time on the LC column, MS1 spectral information using a ±5 ppm mass accuracy range, and characteristic MS2 fragments. A library of retention time, MS1, and MS2 information under our LC-MS/MS conditions was generated using purified compounds from commercial sources, including the Metlin-tested mass spectrometry metabolite library (MSMLS) from Sigma-Aldrich.

Each metabolite from the Mass Spectrometry Metabolite Library (MSMLS, Sigma-Aldrich) listed in Figure S2 was directly infused into the mass spectrometer, and the optimum conditions for ESI and MS/MS fragmentation were defined. Metabolites were used to determine the ionization and MS settings used in the untargeted metabolomics studies. Further, these metabolite standards were used to build a MS2 fragment library used for metabolite identification. Kynurenine standard (Sigma-Aldrich K8625) was used to generate retention data listed in Figure 2C.

### Small Molecule Inhibitor and Activator Treatment

NLG919 (Cayman Chemicals #21509) was used to inhibit IDO1 activity. CH223191 (Tocris Bioscience #3858) was used to inhibit AhR, and 2,3,7,8-tetrachlorodibenzo-p-dioxin (TCDD; Cambridge Isotopes Laboratories ED-901) was used to activate AhR. NLG919 and CH223191 were suspended in DMSO (Sigma-Aldrich D2650). Cells treated with DMSO were used as a control for NLG919 and CH223191. Since TCDD is a neat compound (e.g., liquid at room temperature), molecular biology grade nuclease-free sterile water (Grow Cells NUPW-0500) was used as a control. For all experiments using small molecules, the compound was added to cells after infection. At 2 hpi, cells were washed with PBS. Cells were then fed growth medium that contained either 100nM NLG919, 3μM CH223191, 3.1μM TCDD, DMSO, or water. At 3 and 6 dpi, the growth medium was removed and replaced with medium containing fresh compounds. Cell viability under small molecule treatment conditions was determined using a Lactate Dehydrogenase (LDH) release Cytotoxicity Assay kit (Thermo Pierce 88953).

### Reverse-Transcription Quantitative PCR (RT-qPCR)

RT-qPCR was performed to determine the level of mRNA expressed in NT and HIF1a KO cells. RNA was isolated and purified from cells using a ZR-Duet DNA/RNA MiniPrep kit (Zymo Research). Next, 1 μg of RNA was converted to cDNA using an oligo dT primer and the Transcriptor First Strand cDNA Synthesis kit (Roche Molecular Systems). qPCR was performed using PowerUp SYBR Green (Applied Biosystems Thermo Fisher Scientific) on an ABI real-time qPCR 7300 instrument. H6PD was used to normalize all samples. The following primer pairs were in this study: IDO1 pair 1 (5’-GGCTTTGCTCTGCCAAATCC-3’ / 5’-TTCTCAACTCTTTCTCGAAGCTG-3’), IDO1 pair 2 (5’-GCATTTTTCAGTGTTCTTCGCATA-3’ / 5’-CATACACCAGACCGTCTGATAGCT-3’), VEGF (5’-TCCTCACACCATTGAAACCA-3’ / 5’-GATCCTGCCCTGTCTCTCTG-3’), and H6PD (5’-GGACCATTACTTAGGCAAGCA / 5’-CACGGTCTCTTTCATGATGATCT-3’).

### KYN Treatment and Visualization of HCMV Plaque

KYN (Sigma-Aldrich K8625) was suspended in 0.1N HCl. A 1 mM concentration of KYN was added to growth medium (RPMI 1640, 10% FBS, 5mM HEPES pH 7.4, pen/strep), and the pH was adjusted to pH 7.4 using 0.2 N NaOH. As a control, the same volume of 0.1 N HCl that lacked KYN and 0.2 N NaOH was added to the same volume of growth medium to generate the 0 mM KYN control growth medium. The 1 mM KYN growth medium was serially diluted with the 0 mM KYN growth medium to generate 0.5 mM and 0.25 mM KYN growth medium stocks. All growth medium was sterilized by filtration. At 1 dpi, HFF cells were seeded onto a 6-well plate at a concentration of 1.6×10^5^ cells per well. Cells were infected with 100 infectious units per well with TB40/E for 1 h. After infection, the cells were washed with PBS and 2 mL of growth medium containing 0, 0.25, 0.5, or 1 mM KYN and 0.6% methylcellulose was added. At 3 dpi, the growth medium was removed and replaced with fresh growth medium to replenish the levels of KYN. At 6 dpi, the cells were fixed with methanol. The IE1-positive nuclei were visualized using immunofluorescence using an antibody against IE1 as previously described (5). The IE1-positive nuclei were counted in at least 10 plaques per well with a plaque being defined as having three or more IE1-positive nuclei. All nuclei were visualized using Hoechst 33342.

## QUANTIFICATION AND STATISTICAL ANALYSIS

Unless otherwise specified, data are displayed as mean ± standard deviation where N equals the number of independent biological experiments. For untargeted metabolomics, statistical analysis was performed in consultation with the Statistical Consulting Lab at the BIO5 Institute, University of Arizona. The untargeted LC-MS/MS data used in this study formed 4 distinct datasets: *i)* intracellular metabolites/positive ion acquisition, *ii)* intracellular metabolites/negative ion acquisition, *iii)* extracellular metabolites/positive ion acquisition, and *iv)* extracellular metabolites/negative ion acquisition. Each of these four datasets was analyzed separately by importing them into R, and log-transformed to approximate normality. Only features present in both HCMV-infected NT cells and HCMV-infected HIF1α KO cells were analyzed, so that imputation of missing datapoints between NT and HIF1α KO conditions was unnecessary. Linear mixed-effect models were used to model the log-transformed measurements, with biological and technical replicates treated as random effects. The model outputs for each feature were a fold change (HCMV-infected HIF1α KO cells relative to HCMV-infected NT cells) and p-value, which were adjusted for multiple comparisons using the Benjamini-Hochberg procedure (49). These analyses were used to generate the volcano plots in Figure 2B. OriginPro 2019 was used for all other statistical analysis.

## Acknowledgments

We thank Dr. Dean Billheimer and Shripad Sinari of the Statistics Consulting Lab at the University of Arizona for assisting with statistical analysis of untargeted metabolomics data. We are grateful to Dr. James Alwine for helpful comments related to this project. Finally, we are grateful to Ian Kline for providing critical feedback. This project was supported by startup funds from the University of Arizona Health Sciences, College of Medicine-Tucson, and BIO5 Institute to J.G.P. Additional support for this work was provided by a New Investigator Award (ADHS18-198868) to J.G.P. from the Arizona Biomedical Research Commission, made available through the Arizona Department of Health Services.

## Author Contributions

L.M.W., Y.X., and J.G.P. conceived, designed, and performed the experiments; all authors assisted with analyzing the data, interpreting the results, and writing of the manuscript.

## Supplemental Figures

**Figure S1.**
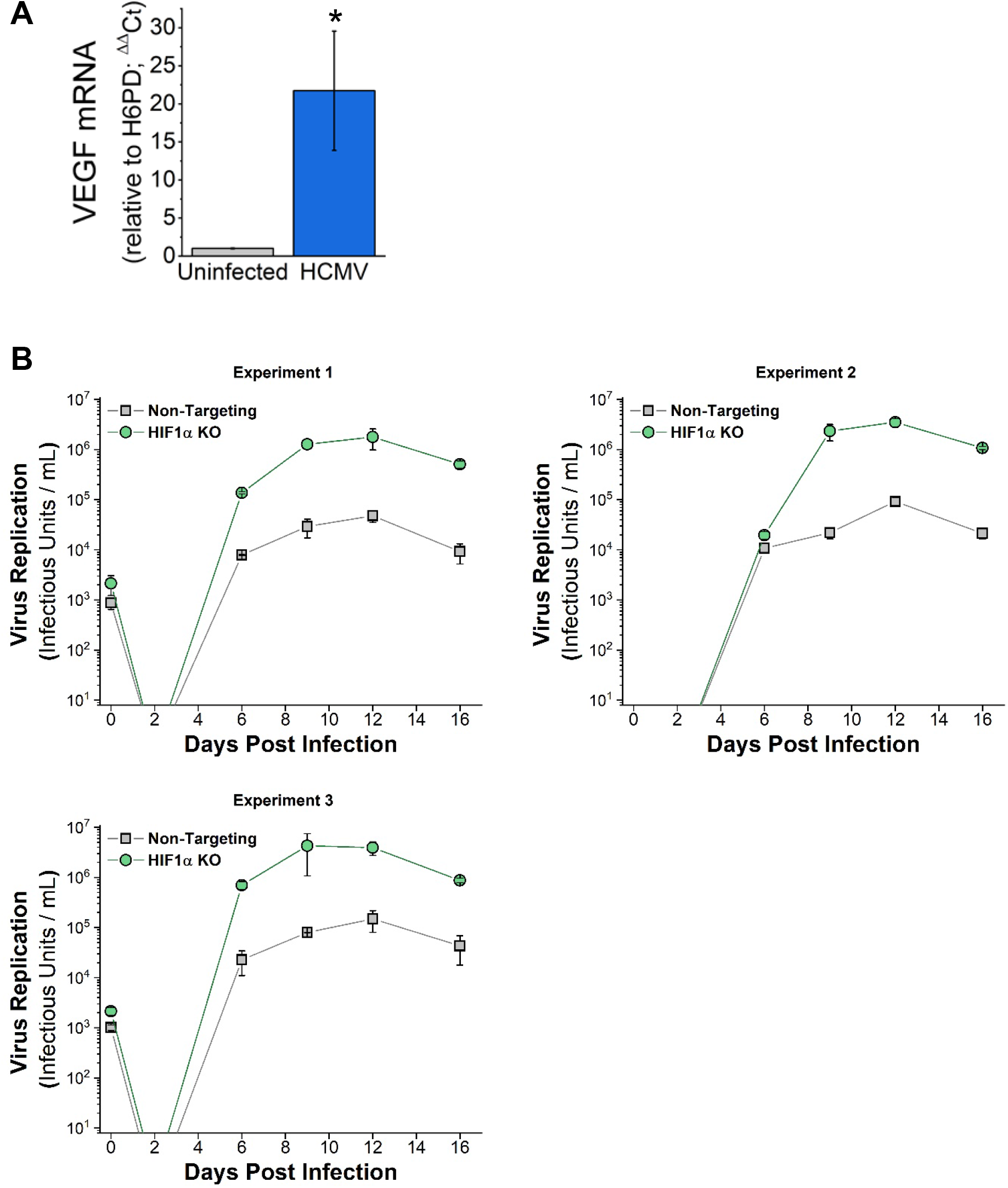
HCMV replication in NT and HIF1α KO cells. **(A)** VEGF transcripts were quantified at 2 dpi using reverse transcriptase quantitative PCR (RT-qPCR) relative to the housekeeping gene H6PD. Fibroblast cells were mock-infected or HCMV-infected at MOI=3 in normoxia, as described in Figure 1A-B. VEGF transcript levels are represented as mean ± S.D. relative to mock-infected. Significance was determined using an unpaired t test, N=3 (*p<0.05). **(B)** Shown are the individual growth curves for the three independent experiments described in Figure 1E. At every time point, two independent TCID_50_ assays were performed for each sample. The error bars represent the S.D. of the two TCID_50_ assays.

**Figure S2.**
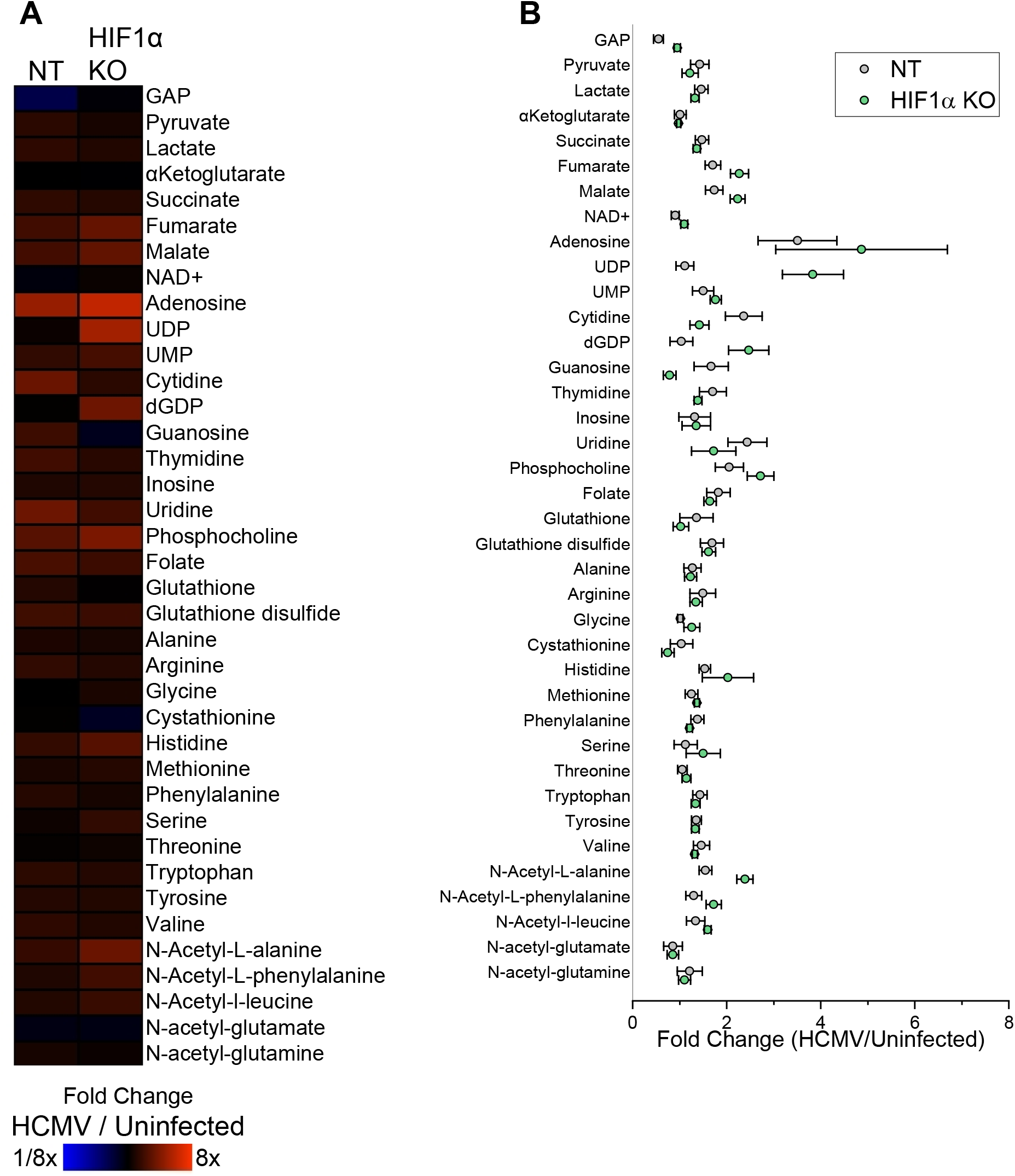
Metabolomic analysis of NT and HIF1α KO cells. **(A)** Metabolites in HCMV-infected NT and HIF1α KO cells were measured at 2 dpi. Metabolites were identified and quantified using LC-MS/MS. The data is represented as fold change of HCMV-infected NT relative to uninfected NT cells and HCMV-infected HIF1α KO relative to uninfected HIF1α KO cells. The conditions are described in Figure 2. **(B)** The data in part A are represented as the mean ± S.E.M.

**Figure S3.**
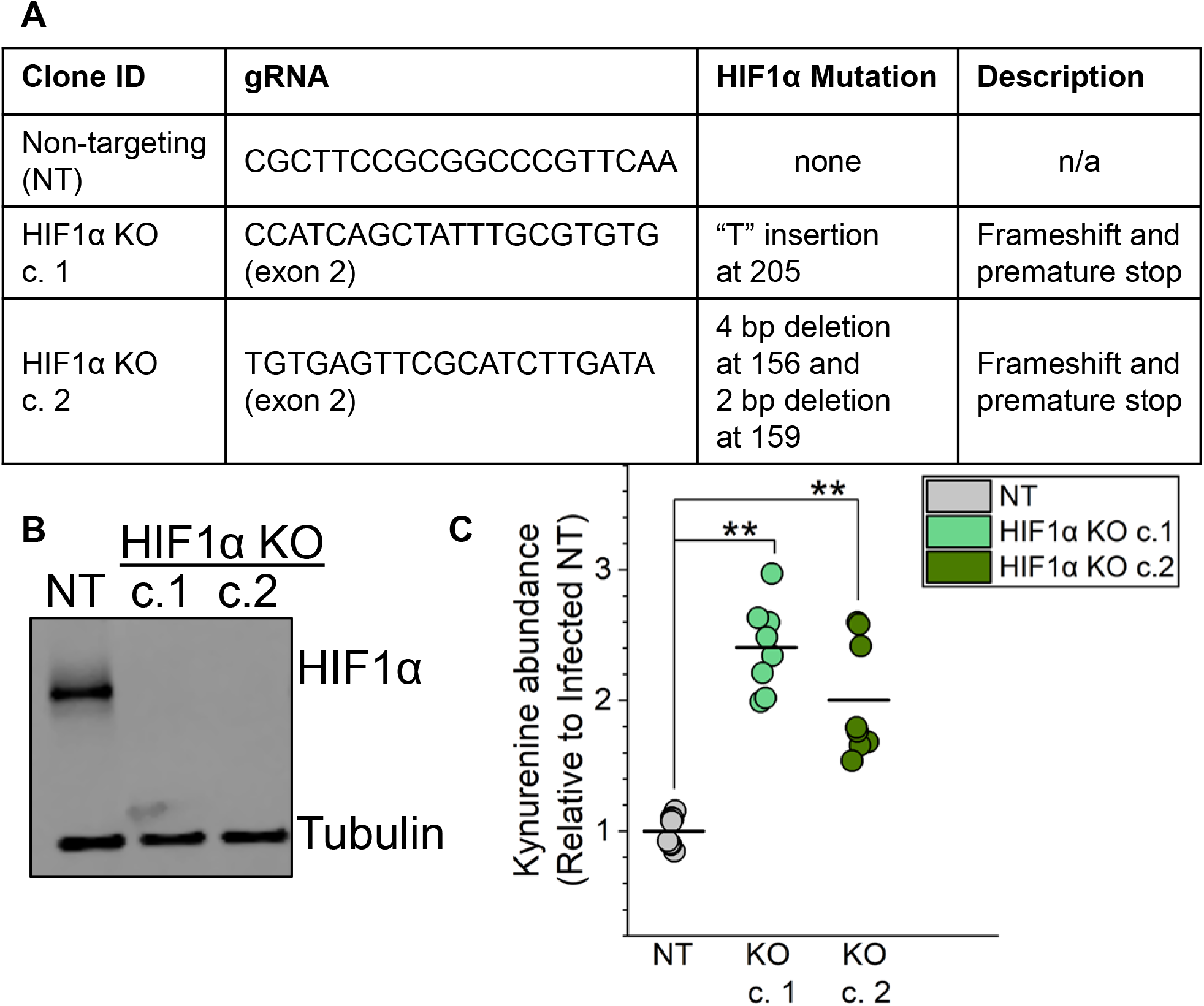
Characterization of two HIF1α KO clones. **(A)** Description of CRISPR/Cas9 gRNAs used in this study. Two independent HIF1α KO clones were generated and confirmed by Indel sequencing. The mutations identified and their consequences on HIF1α protein are shown. **(B)** HIF1α protein levels in NT and two HIF1α KO clones following 24 h treatment with 100 μM CoCl2 to mimic hypoxia. **(C)** KYN levels in HCMV-infected NT and HIF1α KO clones 1 and 2. Cells were infected at MOI=2 with TB40/E. At 2 dpi, metabolites were extracted and KYN was measured by LC-MS/MS. The means are represented by a black bar. Four independent experiments were performed, each with duplicate samples per condition. Each of the 8 samples measured are represented by a dot. The data is represented as fold change relative to HCMV-infected NT cells, and significance was determined by two-way ANOVA, Tukey test, N=4 (**p<0.01).

**Figure S4.**
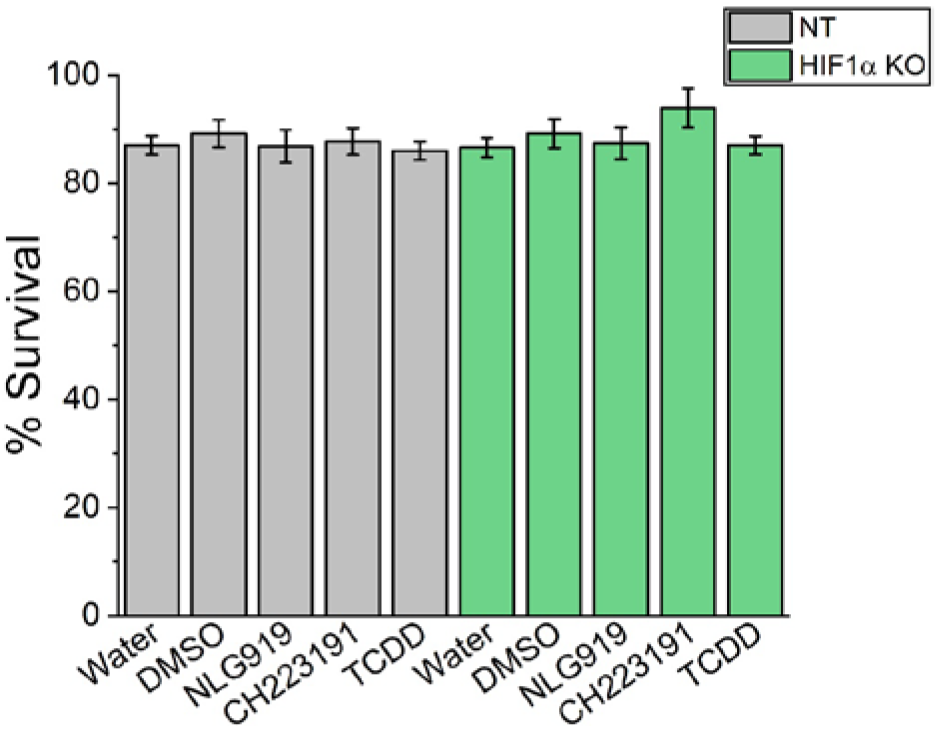
Survival of Cells treated with NLG919, CH223191, and TCDD. Cytotoxicity of uninfected NT and HIF1α KO cells treated with 100nM NLG919, 3μM CH223191, or DMSO was determined. Similarly, cells were treated with 3.1nM TCDD or water as a control. Cell survival was measured at 9 days post treatment using a LDH Cytotoxicity kit. N=4. Data are represented as mean ± S.E.M.

